# CRISPR/Cas9-mediated generation of two isogenic *CEP290*-mutated iPSC lines

**DOI:** 10.1101/2025.03.31.646311

**Authors:** Joana Figueiro-Silva, Melanie Eschment, Michelle Mennel, Affef Abidi, Beatrice Oneda, Anita Rauch, Ruxandra Bachmann-Gagescu

## Abstract

*CEP290* is an important human disease gene, as mutations are implicated in a broad spectrum of autosomal recessive ciliopathies, including Leber congenital amaurosis and Joubert, Meckel, Senior-LØken or Bardet Biedl syndromes. To create isogenic mutant human induced pluripotent stem cell (hiPSC) lines for disease modeling, we employed CRISPR/Cas9 to introduce disease-relevant mutations into the control hiPSC line HMGU1 (ISFi001-A). Thorough characterization of the lines, including the effect of the mutation at the mRNA and protein level, shows that these *CEP290*-mutant lines provide a useful resource for studying ciliopathy disease mechanisms and cilia biology through differentiation into diverse cell types and organoids.

## Resource Table

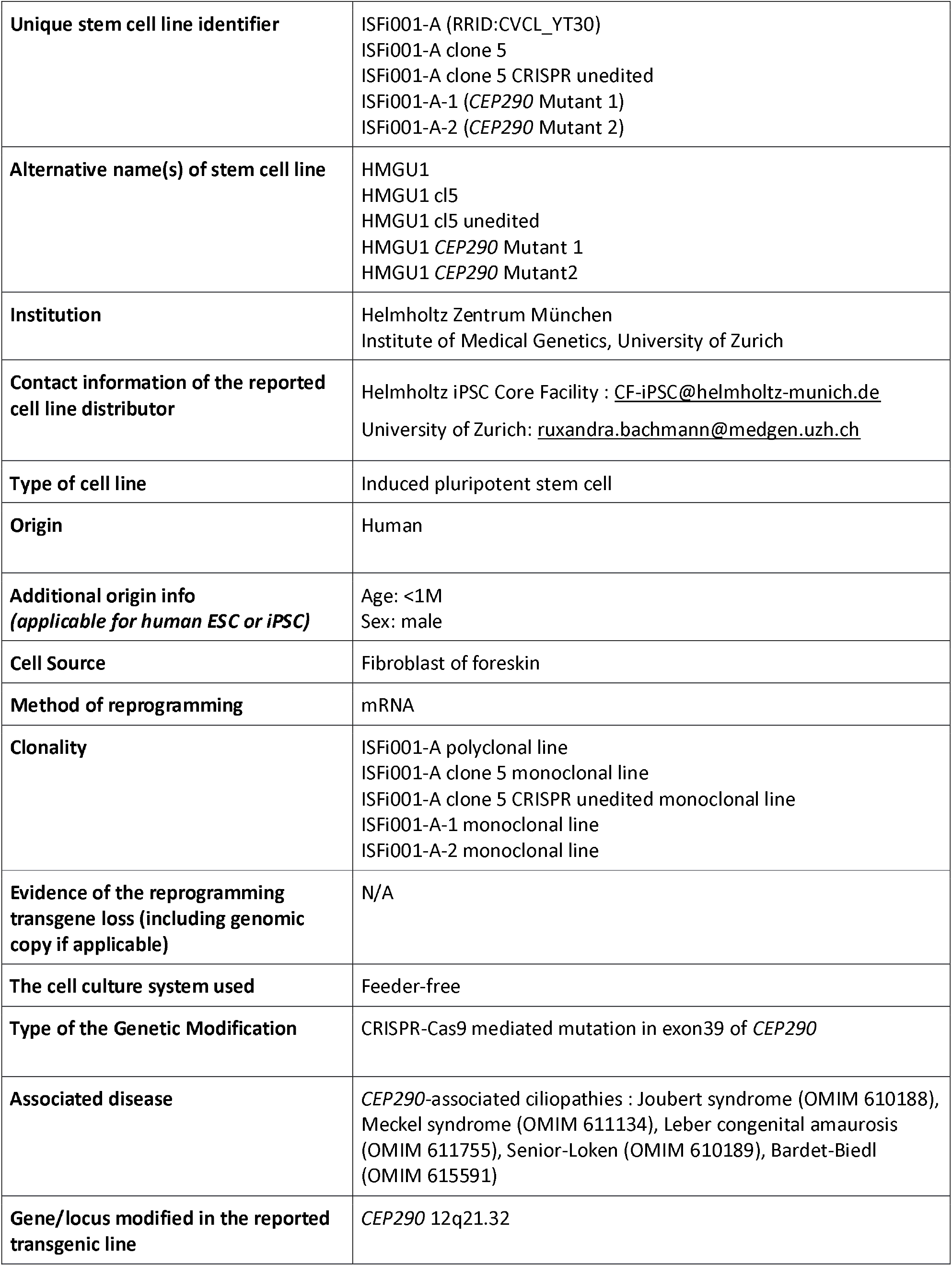
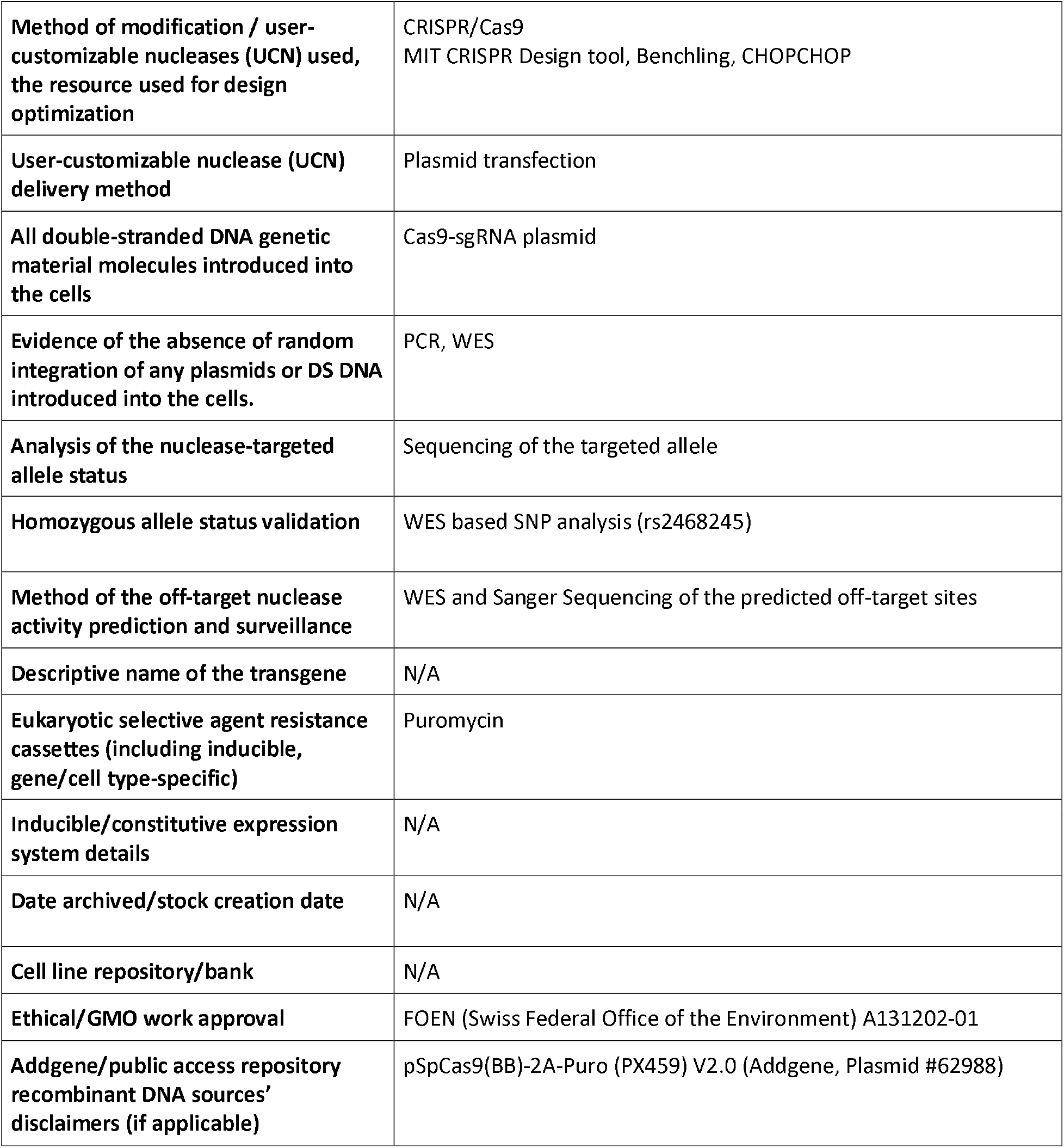

## Resource utility

This study utilized CRISPR/Cas9 to engineer isogenic *CEP290*-mutant hiPSC lines, providing a valuable resource for investigating ciliopathies which are important monogenic disorders. These cell lines enable disease modeling and cilia biology research through differentiation into various cell and organoid types, offering insights into the molecular mechanisms of *CEP290*-related disorders.

## Resource Details

*CEP290* is an important human disease gene, mutated in several syndromic and non-syndromic ciliopathies (Coppieters et al., 2010; Drivas and Bennett, 2014; Bachmann-Gagescu et al., 2015; Reiter and Leroux, 2017). The functional diversity of cilia across cell types (Mill et al., 2023) requires models that reflect tissue-specific ciliary biology and disease mechanisms, which can be achieved with iPSC-derived models. We employed CRISPR-Cas9 genome editing to introduce mutations in *CEP290* into the previously established healthy control polyclonal hiPSC line HMGU1 (ISFi001-A) which we first subcloned (HMGU1 cl.5 / ISFi001-A cl.5). Given clustering of disease mutations in the C-terminal myosin-tail homology domain of *CEP290*, we designed two CRISPR-Cas9 single guide RNAs (sgRNAs) to target this region, specifically exon 39 (Figure 1A). Both sgRNAs were inserted into a non-integrating plasmid. We chose two mutant clones (*CEP290* Mutant 1 / ISFi001-A-1 and *CEP290* Mutant 2 / ISFi001-A-2), and one clone that was not edited at the target site after CRISPR’ing as a control line (HMGU1 cl5 unedited / ISFi001-A cl.5 unedited). Sanger sequencing demonstrated Mutant 1 to harbor a homozygous 5-basepair deletion at target site 1 resulting in a frameshift with premature stop codon (p.Leu1749Serfs*10) and predicted to cause nonsense mediated mRNA decay (NMD). Mutant 2 displayed a homozygous 5-basepair substitution at the start of exon 39 and a homozygous 11-basepair deletion (4 intronic and 7 exonic bps) at target site 2 predicted to result in complete loss of the splice donor site. Using end-point RT-PCR targeting exons 36 to 41, we identified additional bands on gel electrophoresis in both mutants compared to controls (Figure 1B). Sanger sequencing after band excision confirmed that the shorter band found in both mutants corresponds to skipped exon 39, while a slightly larger than full-length band in Mutant 2 corresponds to 18bp retained from intron 39-40 in addition to the 7bp exonic deletion at target site 2 (Figure 1B). Prediction tools indicate that the combined effect of the 7 exonic bp deletion with the 18 bp intronic retention leads to a frameshift with premature stop codon (p.Leu1787Hisfs*12). Since there is incomplete NMD, we quantified residual mRNA amounts using RT-qPCR analysis, demonstrating ∼62% reduction in *CEP290* total mRNA levels in Mutant 1 and ∼75% reduction in Mutant 2 (Figure 1C). We next assessed *CEP290* protein levels using SDS-page with an antibody targeting the C-terminus of *CEP290*. This showed minimal residual protein, likely lacking the in-*frame* exon 39, in both mutants, visible only when loading high protein amounts (Figure 1D). Since nonsense-induced alternative splicing of in *frame* exons has been described for *CEP290* in patients with ciliopathies (Drivas et al., 2015), the generated lines reproduce the disease situation. We next assessed pluripotency of the hiPSC lines by immunofluorescence staining OCT3/4 and Nanog (Figure 1F) and by RT-qPCR for *OCT3/4, NANOG* and *SOX2* (Figure 1G), confirming comparable expression in all cell lines. We confirmed the hiPSC lines’ ability to differentiate into all three germ layers in vitro, with expression of lineage markers: HAND1, SM22a for mesoderm, *SOX17* and *FOXA2* for endoderm and *PAX6* and *MAP2* for ectoderm (Figure 1H). A SNP-array analysis ruled out major chromosomal aberrations (Figure 1I) and Sanger and whole exome sequencing ruled out off-target sequence variants resulting from editing and passaging.

**Figure 1:**
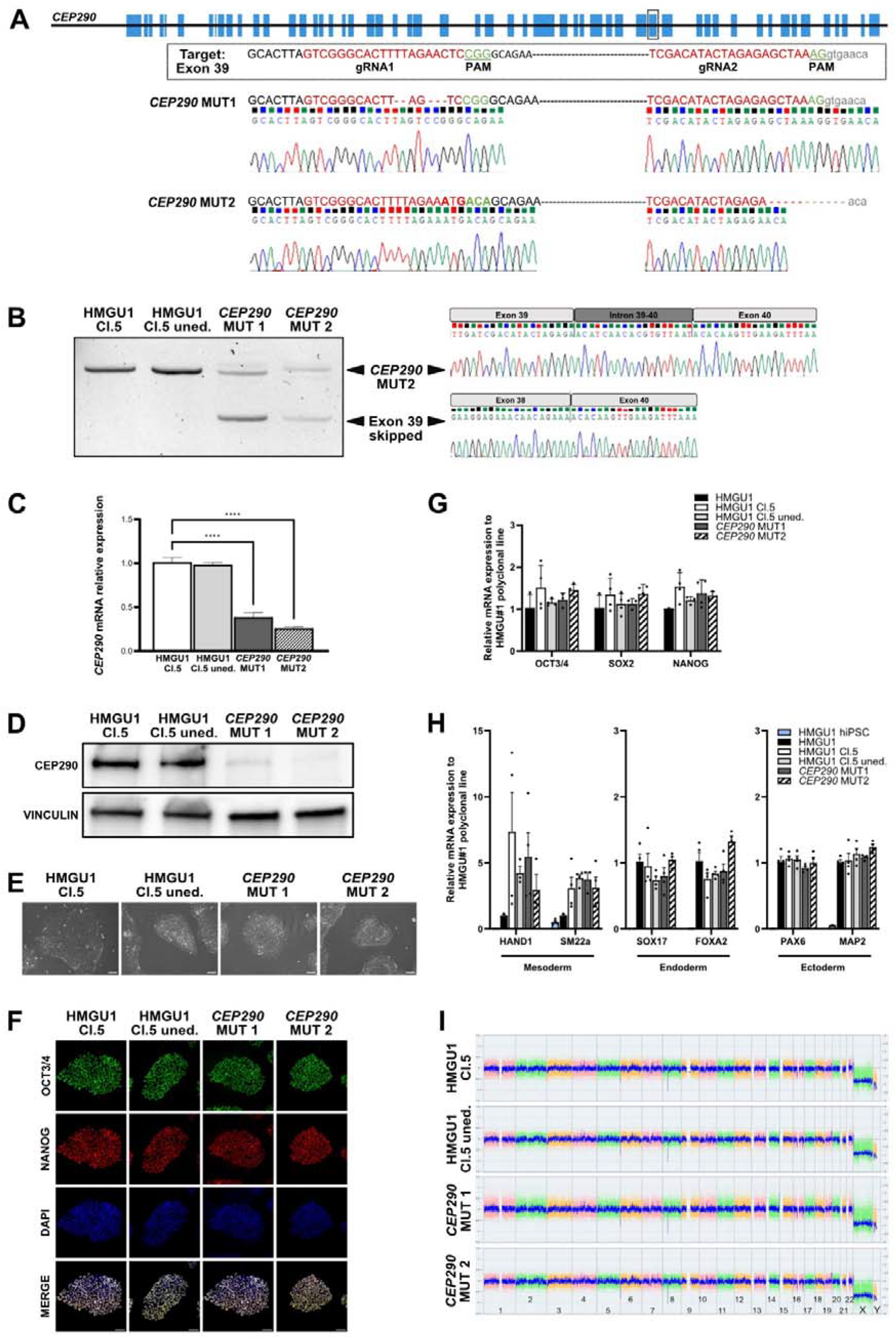
Generation, characterization and quality control of *CEP290*-iPSC lines.

## Materials and Methods

### Cell culture and generation of CEP290 mutant human iPSC lines

The polyclonal HMGU1/ISFi001-A line was subcloned and all subsequent experiments used clone 5 (ISFi001-A cl.5). hiPSCs were maintained in StemMACS iPS-Brew XF (Miltenyi Biotec) media on Geltrex™ LDEV-free coated vessels at 37°C and 5% CO_2_. Colonies were EDTA (0.5mM) passaged when reaching 70-80% confluence. Single guide RNAs (sgRNAs) for targeting *CEP290* were cloned into pSpCas9(BB)-2A-Puro (PX459) V2.0 (Addgene Plasmid #62988) and transformed in One Shot™ Stbl3™ Chemically Competent E. coli (Thermofisher). Cells were electroporated using the Amaxa 4D nucleofector (program CA137, Lonza) and P3 Primary Cell Kit (Lonza). 48h after electroporation, cells were treated with puromycin (0.4 ug/mL). Single cell dilution was performed and drug-resistant clones were picked manually, expanded and subsequently sequenced.

### Genomic DNA extraction, PCR and Sanger Sequencing

Genotyping was performed by genomic DNA (gDNA) extraction using QuickExtract DNA Extraction Solution (LGC), standard PCR and Sanger Sequencing. Alamut visual plus tool (Sophia Genetics) was used to predict the effect of genomic mutations.

### Genomic integrity QC with SNP array and Whole Exome Sequencing (WES)

For SNP array and WES analysis, gDNA of cells at passages 34-38 was extracted using the DNeasy Blood & Tissue Kit (Qiagen). For SNP-array, gDNA was analyzed with the CytoScan HD-array (Affymetrix Inc) at a genome-wide resolution of 200kb for copy number variants according to manufacturer’s instructions. Analysis was performed with the Chromosome Analysis suite (ChAS) software (Affymetrix). WES was performed using the Twist Exome 2.0 plus Comprehensive Exome Spike-In and the Twist Mitochondrial Panel. After paired-end sequencing (NovaSeq 6000 S2 Reagent Kit (318 cycles), 159 fwd-159 rev) on a NovaSeq 6000 sequencer (Illumina Inc.), the coding regions as well as the exon-intron-boundaries were analyzed up to 20 base pairs into the intronic regions. Alignment was performed with DRAGEN Bio-IT Platform v4.3 (Illumina Inc.) and analyzed using VIA v7.1 (Bionano Genomics, Inc.). In both Sanger and WES sequencing we compared the results of all edited clones to those of the isogenic control hiPSC line ISFi001-A cl.5.

### Reverse Transcription-Quantitative PCR (RT-qPCR)

Total RNA was extracted using the RNeasy Plus Mini Kit (Qiagen) and reverse transcribed using the High-Capacity RNA-to-cDNA Kit (Applied Biosystems). Gene expression was assessed by qPCR using PowerUp SYBR Green Master Mix (Applied Biosystems). The polyclonal ISFi001-A hiPSC line was used as reference. Samples were normalized to housekeeping genes GAPDH and CAPN10. Cells were tested at passages 32 to 38.

### Western Blot

Cells were lysed using RIPA lysis buffer and SDS-Page was performed following standard procedures, using the antibodies listed in table 2.

**Table 1:**

| Classification | Output type | Result | Data |
| --- | --- | --- | --- |
| **Schematic of a transgene/genetic modification** | Schematic illustrating the structure and location of the introduced genetic modification | Visual representation of gene editing at exon 39 | Fig. 1 panel A |
| **Morphology** | Photography | Normal Morphology | Fig. 1 panel E |
| **Pluripotency status evidence for the described cell line** | Qualitative analysis (*Immunocytochemistry*) | All lines are positive for OCT3/4 and NANOG | Fig. 1 panel F |
|  | Quantitative analysis (*RT-qPCR*) | All lines show strong expression of *OCT3/4*, *SOX2* and *NANOG* | Fig. 1 panel G |
| **Karyotype** | SNP array | 46,XY, Resolution: 200kb | Fig. 1 panel I |
| **Genotyping for the desired genomic alteration/allelic status of the gene of interest** | PCR across the edited site or targeted allele-specific PCR | PCR followed by Sanger sequencing | Fig. 1 panel A |
|  | Evaluation of the - (homo-/hetero-/hemi-) zygous status of introduced genomic alteration(s) | Sanger Sequencing analysis including the adjacent SNP rs2468245 | Supplementary Fig.1 panel A |
|  | Transgene-specific PCR (when applicable) | N/A |  |
| **Verification of the absence of random plasmid integration events** | PCR/Southern | N/A |  |
| **Parental and modified cell line genetic identity evidence** | WES analysis of SNP panel for genetic identity evidence | Identical SNPs for all lines | Supplementary Table |
| **Mutagenesis / genetic modification outcome analysis** | Sequencing (genomic DNA PCR or RT-PCR product) | Sanger Sequencing of region of interest:<br><u>ISFi001-A-1:</u><br>NM_025114.4(CEP290):c.5245_5251delinsGA<br>p.(Leu1749Aspfs*10) hg19<br>chr12:g.88472982_88472988delinsTC, homozygous<br><u>ISFi001-A-2:</u><br>NM_025114.4(CEP290):<br>c.5251_5256delinsACAGTA(;<br>);5358_5364+4del<br>p.(Leu1751_Arg1752delinsThrVal)(;)? hg19<br>chr12:g.88472977_88472982delinsTACTGT(;);88472867_88472877del,<br>homozygous | Fig. 1 panel A |
|  | PCR-based analyses | **RT-PCR and qPCR:**<br>Both mutations lead to incomplete NMD, with 38% (Mutant 1) and 25% (Mutant 2) residual mRNA. Band including skipped exon 39 observed. | Fig. 1 panel B |
|  | Western blotting (for knock-outs, KOs) | Minimal residual CEP290 protein levels in both mutant lines. | Fig. 1 panel D |
| **Off-target nuclease activity analysis** | PCR across top 5/10 predicted top likely off-target sites, whole genome/exome sequencing | No mutations at predicted-off-target sites by Sanger sequencing and WES. | Supplementary Fig.1 panel B |
| **Specific pathogen-free status** | Mycoplasma testing by endpoint PCR | Negative | Supplementary Fig.2 |
| **Multilineage differentiation potential** | Directed differentiation | All lines differentiate into all three germ layers | Fig. 1 panel H |
| **List of recommended germ layer markers** | Expression of markers at mRNA level (RT-qPCR) | Ectoderm: *PAX6*, *MAP2*<br>Endoderm: *SOX17*, *FOXA2*<br>Mesoderm: *SM22a*, *HAND1* | Fig. 1 panel H |
| ***Outcomes of gene editing experiment (OPTIONAL)*** | Brief description of the outcomes in terms of clones generated/establishment approach/screening outcomes | See above |  |
| ***Donor screening (OPTIONAL)*** | HIV 1 + 2 Hepatitis B, Hepatitis C | N/A |  |
| ***Genotype - additional histocompatibility info (OPTIONAL)*** | Blood group genotyping | N/A |  |
|  | HLA tissue typing | N/A |  |

**Table 2:**
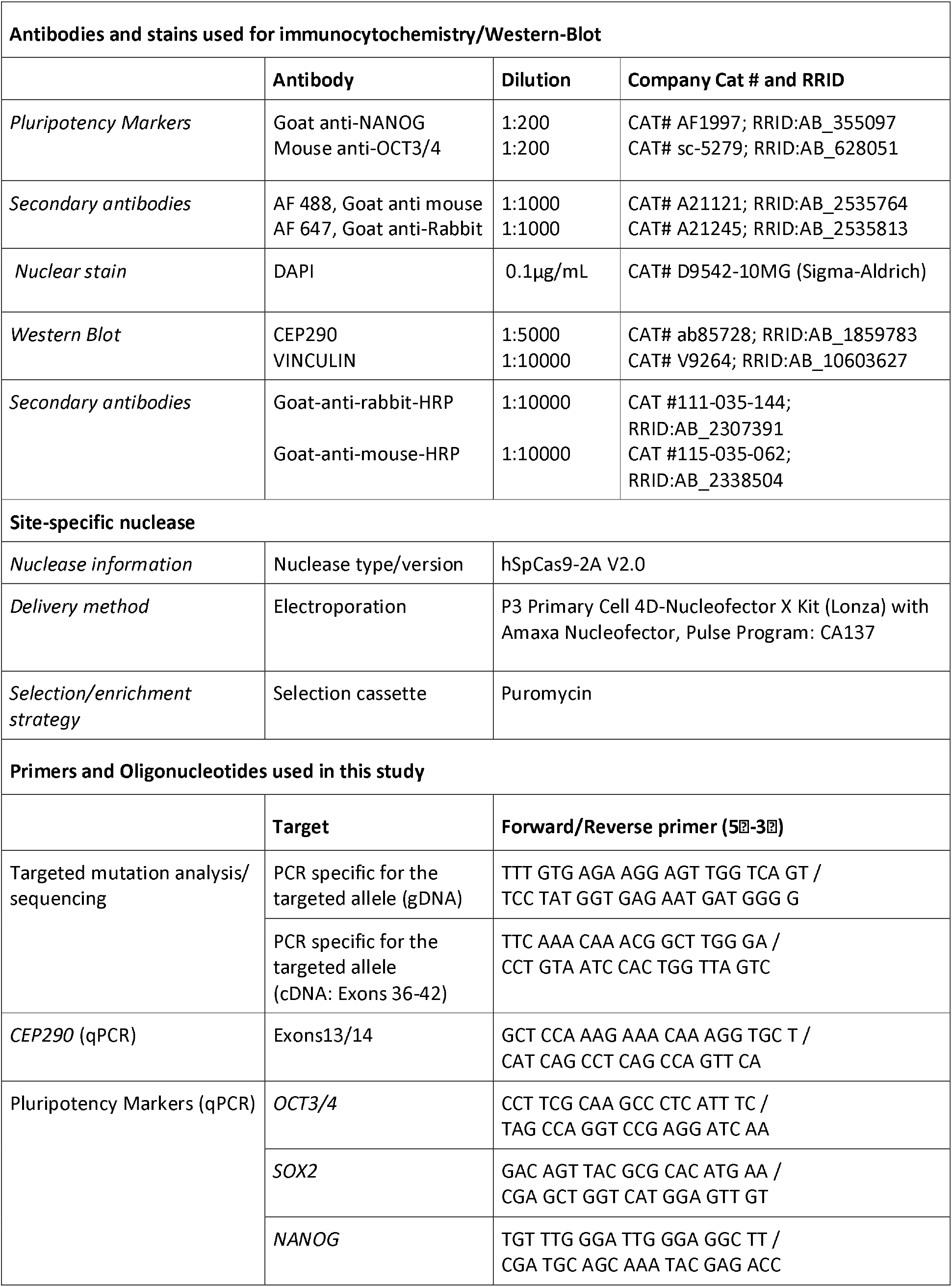
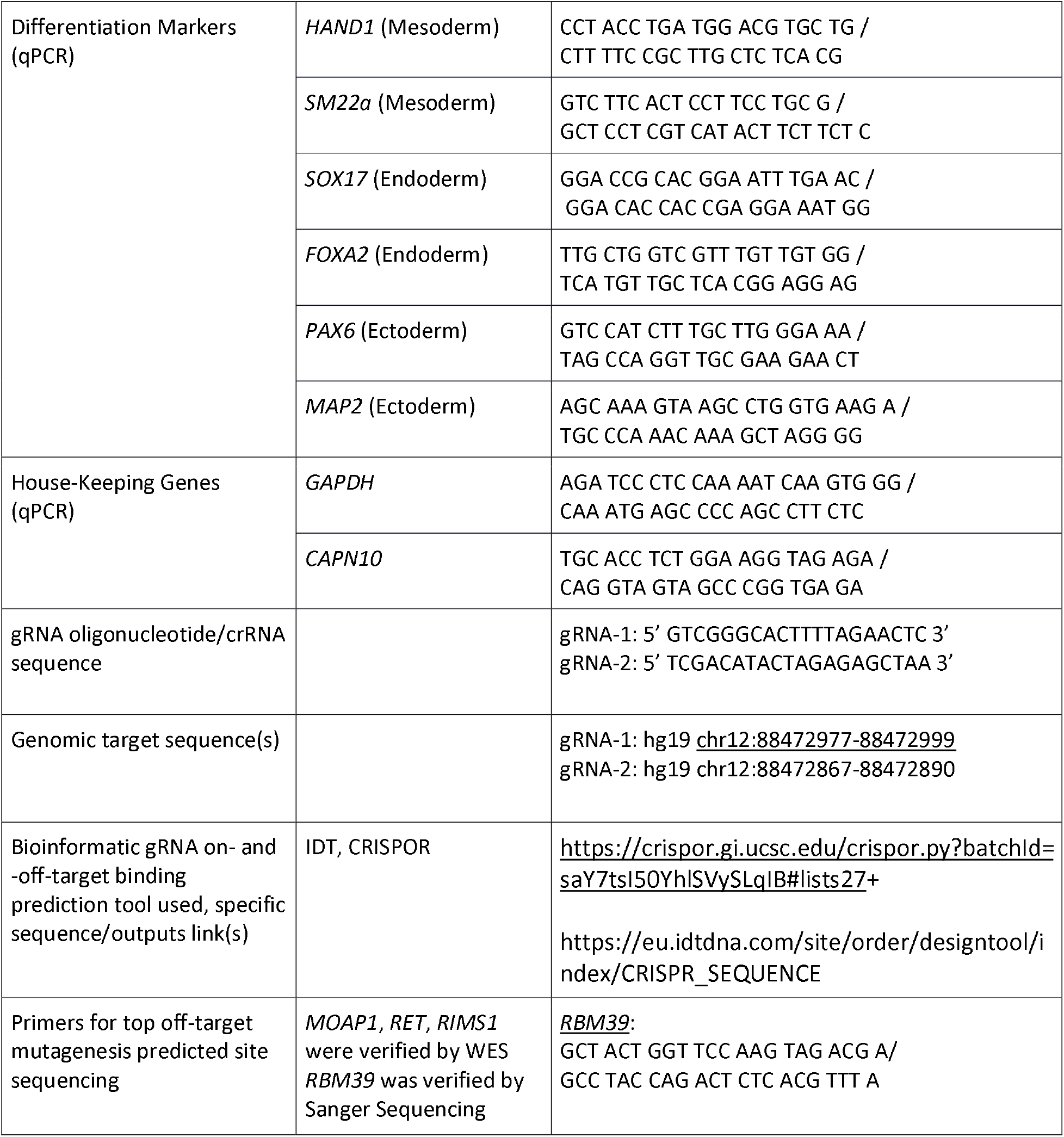

### Immunofluorescence

Cells at passages 30 to 38 were fixed for 10 min in 4% PFA, quenched with PBS-Glycine for 5 min and washed twice with 1X PBS. Fixed specimens were permeabilized and blocked with 1% BSA, 1%NGS, 0.3% Triton-X-100 in PBS for 1 h. Primary antibody staining was done at 4°C overnight and secondary antibody staining at room temperature for 2 h. Nuclei were counterstained for 1h with DAPI. Widefield imaging was done on a Zeiss Axis Scan Z1.

### Differentiation into three germ layers

The differentiation capacity was assessed by directed differentiation to the three germ layers using the StemMACS™ Trilineage Differentiation Kit (Miltenyi Biotec) according to the manufacturer’s protocol.

### Mycoplasma detection

For mycoplasma detection the Venor GeM Classic Kit (Minerva Biolabs) was used, following the manufacturer’s instructions. Cells were tested at passages 30 to 38.

## Acknowledgements

We acknowledge the technical support of the Core Facility iPSC at Helmholtz Zentrum München which provided the HMGU1 hiPSC line. This work was supported by SNSF grant 310030_220012 and the University of Zurich Clinical Research Priority Program *praeclare* and University Research Priority Program *AdaBD*.

## Figures

**Supplementary Figure 1:** Evaluation of homozygous state and Off-target analysis.

**Supplementary Figure 2:** Mycoplasma testing by endpoint PCR.

