## Supplemental data for "CRISPR/Cas9-mediated generation of two isogenic *CEP290*-mutated iPSC lines"

**Supplementary information, Figueiro-Silva et al.
Supplementary Figure 1:**

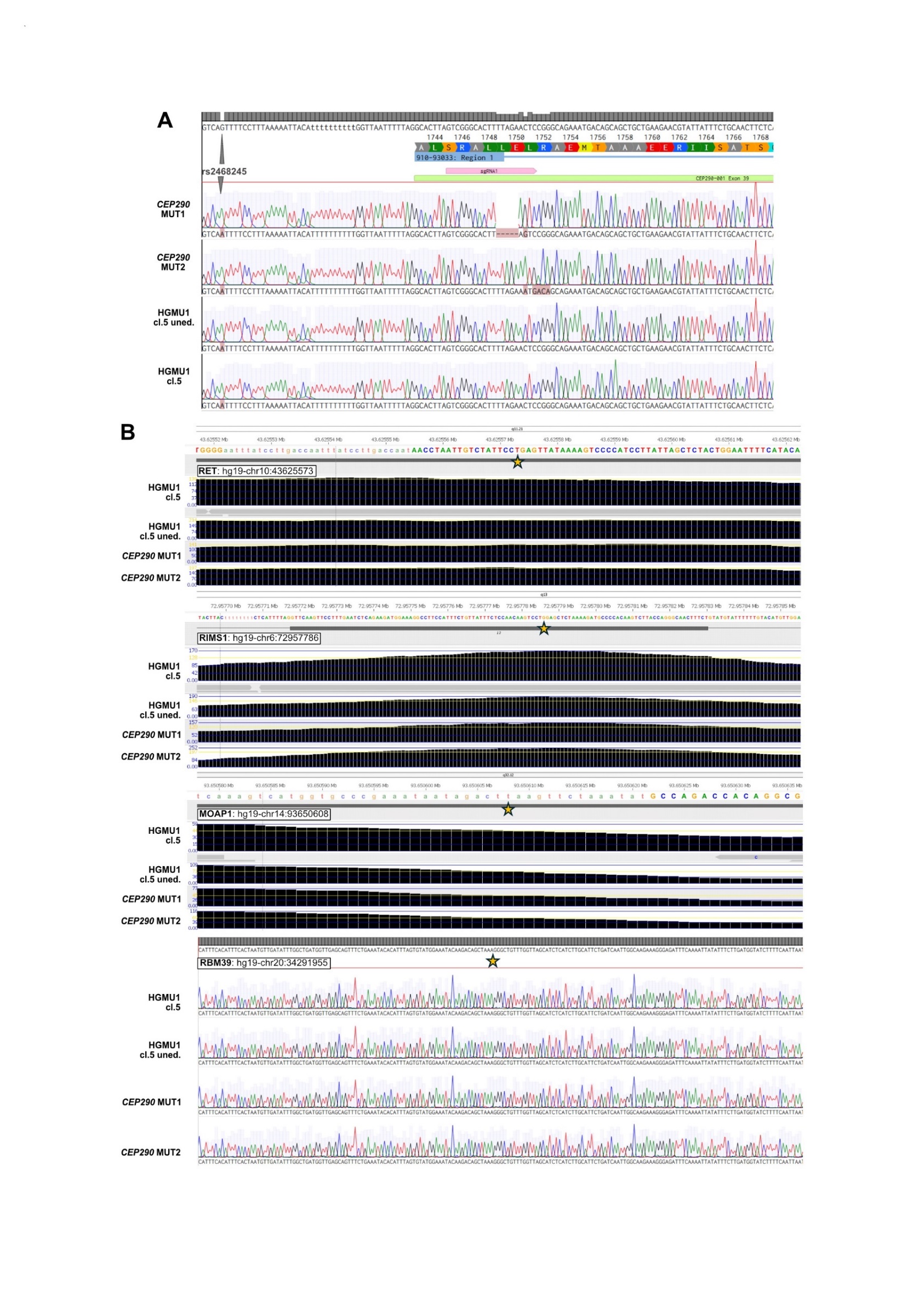

**Supplementary Figure 2:**

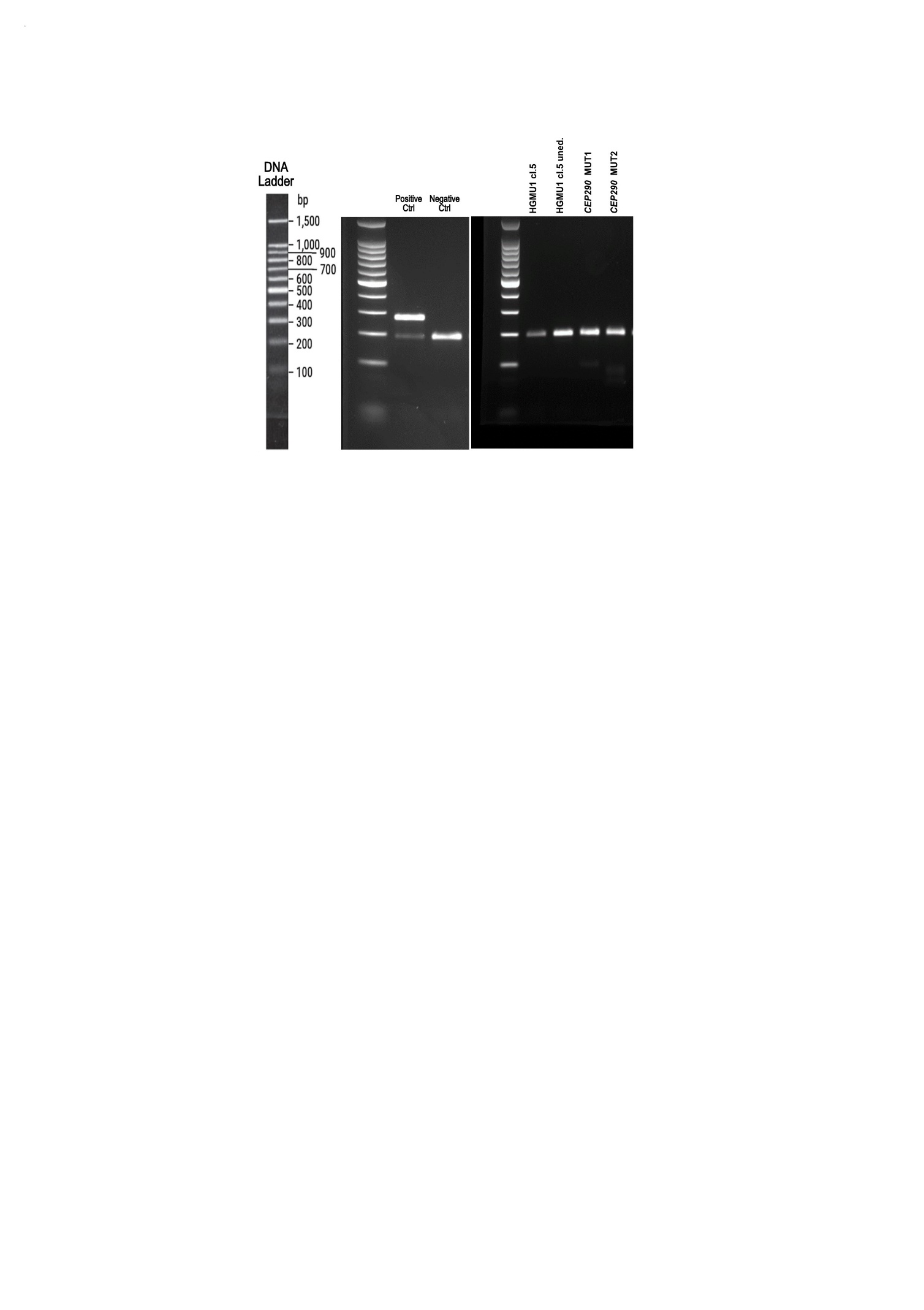

**Supplementary Table: SNP panel for genetic identity evidence.**

|  | **rs12221474** | **rs5960** | **rs6061243** | **rs1128925** | **rs9620123** | **rs8048410** | **rs6503070** | **rs231399** | **rs12990557** | **rs2297079** | **rs12179** | **rs3734557** | **rs1051614** | **rs2279819** | **rs2304035** | **rs6559167** | **rs3741097** | **rs4758686** | **rs3737171** | **rs3744877** | **rs2249057** | **rs3743399** |
| --- | --- | --- | --- | --- | --- | --- | --- | --- | --- | --- | --- | --- | --- | --- | --- | --- | --- | --- | --- | --- | --- | --- |
| Reference alleles | A/C | C/T | C/G | G/T | C/G | A/G | C/T | T/G | G/T | C/G | G/A | A/G | C/G | G/C | A/G | C/A/G | C/G | T/C | G/T | G/A | C/A | G/A |
| HGMU1 cl.5 | A/A | T/T | C/C | G/T | C/C | A/A | C/T | T/G | G/T | C/G | G/G | A/G | T/G | G/C | G/G | C/A | C/G | T/T | T/T | G/A | C/C | G/A |
| HGMU1 cl.5 unedited | A/A | T/T | C/C | G/T | C/C | A/A | C/T | T/G | G/T | C/G | G/G | A/G | T/G | G/C | G/G | C/A | C/G | T/T | T/T | G/A | C/C | G/A |
| CEP290 MUT1 | A/A | T/T | C/C | G/T | C/C | A/A | C/T | T/G | G/T | C/G | G/G | A/G | T/G | G/C | G/G | C/A | C/G | T/T | T/T | G/A | C/C | G/A |
| CEP290 MUT2 | A/A | T/T | C/C | G/T | C/C | A/A | C/T | T/G | G/T | C/G | G/G | A/G | T/G | G/C | G/G | C/A | C/G | T/T | T/T | G/A | C/C | G/A |

Reference : Huang Y et al. Development of a coding SNP panel for tracking the origin of whole-exome sequencing samples. *BMC Genomics,* 25:142 (2024)
